# Brain-based Authentication: Towards A Scalable, Commercial Grade Solution Using Noninvasive Brain Signals

**DOI:** 10.1101/2021.04.09.439244

**Authors:** Ronen Kopito, Aia Haruvi, Noa Brande-Eilat, Shai Kalev, Eitan Kay, Dan Furman

## Abstract

Here we report on a field test where we asked if it is feasible to deliver a scalable, commercial-grade solution for brain-based authentication given currently available head wearables. In this study, forty-nine (49) participants completed multiple sessions in their natural home environment over a single week. Participants used an off-the-shelf brain signal measuring headband to record their own brain activity while completing various tasks. Recording sessions were self-operated by the participants and unsupervised by any expert or technician to simulate real world use cases, while also contrasting common research approaches to this topic that rely on data from controlled laboratory conditions. Although brain signals have a non-stationary, complex nature, when participants watched rapidly presented images, our authentication system was able to successfully construct a unique and robust “brain ID” for each participant. Based on this brain ID, we developed a simplified brain-based authentication method that captures distinguishable information with reliable, commercial-grade performance from participants at their own homes. We conclude that noninvasively measured brain signals are ideal for use in biometric authentication systems, especially in environments where head wearables such as headphones or AR/VR devices are used as these devices offer a natural form factor for capturing participant brain ID continuously.

## 1. Introduction

When a user requests access to a system or a device, an authentication process must confirm whether the identity claim of the user is genuine or whether they are an imposter. A simple example of an authentication method is an alphanumeric password like ‘abc123,’ while a more complex example is a digital fingerprint captured by a smartphone sensor. Effective authentication is critical to security for both consumers and enterprises. Because of the high frequency of use of authentication systems, methods need to be both convenient and secure. That balance - between convenience and security - is a defining performance characteristic of authentication systems. The strongest authentication systems are very secure and, often, very cumbersome to implement and maintain. In contrast, weak authentication methods are very convenient, but have been responsible for countless data breaches because of their equivalent ease of being hacked. The “password chaos” of modern life seems to have reached a boiling point and it is clear that future computing systems need improved methods that both deliver increased security along with an increased convenience that ensures adherence, at the human level of behavior, to security protocols.

Biometric authentication is any method that uses natural occurring information to verify a user’s identity. Many biometric authentication systems have already been developed based on fingerprints, faces, palm veins, irises, voices, gaits, and other metrics. There are significant advantages to using biometrics for authentication, since user experience is convenient and fast, it is non-transferable, and usually the system can reach very high performances (high false rejection rate, and low false acceptance rate) typically without requiring much attention, if any, from the user.

There are disadvantages as well however, since biometrics can be faked or stolen and when that happens, the victim cannot simply replace them to avoid impersonation. Certain biometrics, for example fingerprints or faces, can be easily captured today by cameras remotely without any knowledge from the individual that is being surveilled, and once a face or finger ID is compromised, the remedy is extremely difficult.

Brain-based authentication is the process of verifying an individual’s identity by using their brain signal, and as an approach it offers several distinct advantages over other biometric authentication methods. Since at least the late 1980’s^1^ neuroscientists have observed that noninvasively measured human brain signals carry personally identifying information^2,3^ that differentiates between family members and across a broad population.^4,5^ Brain signals, unlike many other biomarkers are concealed: an invisible signal that is never exposed in daily life. Second, brain signals are dynamic, non-stationary and extremely complex. They are the result of a unique series of brain waves superpositioning in a given brain at any moment, and these waves reflect both personal brain function and anatomy. Taken together, this makes brain signals an ideal candidate for use as a biometric^6,7,8^ method. Indeed, many groups have attempted to build biometric authentication systems based on brain signals.^9,10,11,12,13,14,15,16,17,18,19^ Generally, the process involves a machine-learning classifier to identify if a given brain signal belongs to a genuine identity or to an imposter one (Figure 1).

**Figure 1:**
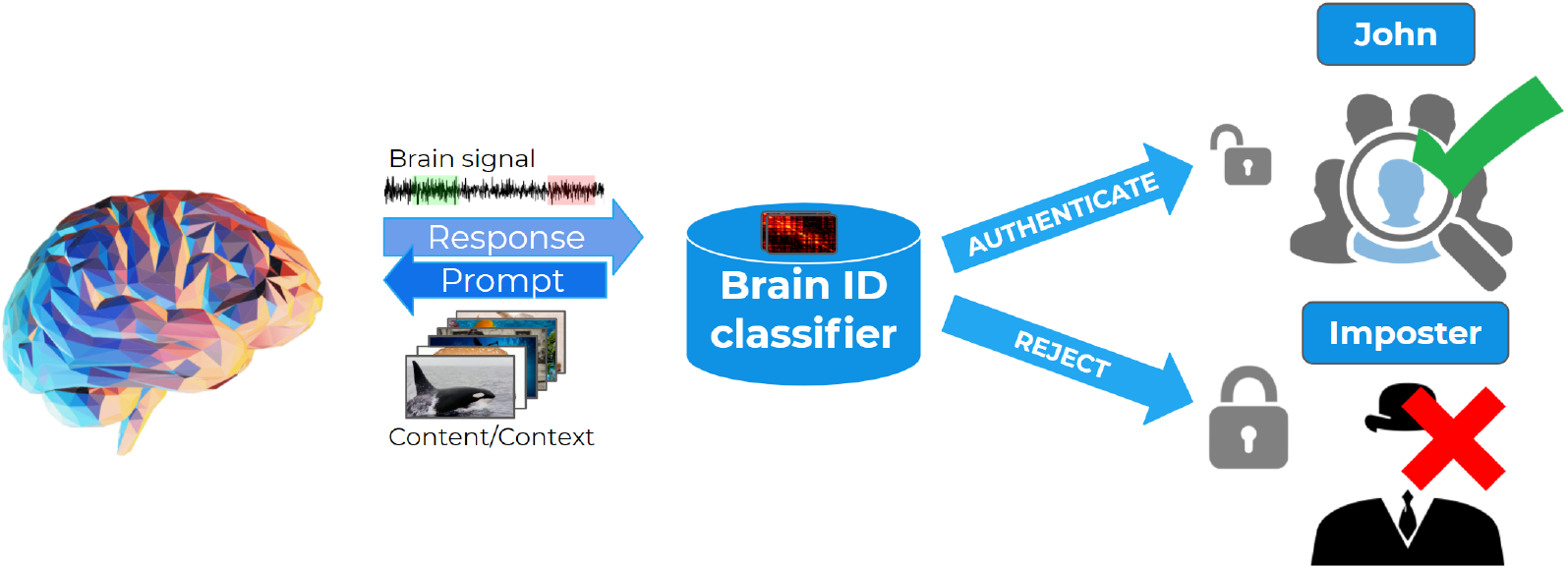
Schematic illustration of the brain-based authentication process. Brain signal is recorded while participants watch images (“Prompt”). The brain response (e.g. “John’s brain signal”) is fed into a trained classifier of that participant. The classifier decides if the brainpattern matches the participant (authenticates “John”) or not (rejects).

The overall usability of brain-based authentication systems has been increasing since 2010,^20^ however most are still far from proving field-viability and bringing new value and utility to existing authentication providers.^21,22^ Brain data is commonly collected in a laboratory under controlled conditions in other studies, where a trained technician is an essential part of the brain measurement procedure. Often authentication performance tests are also done with a small number of people, with all measurements taking place in a single session despite the common industry knowledge that single session data maps poorly to daily consumer electronics usage, where individuals put devices on and off regularly.^23,24,25,26^

While brain biometric identity appears to be one of the most natural, powerful methods for head wearables, its robustness has not been sufficiently vetted in real world conditions that parallel the end use cases such as:

- Professionals who work remotely and wear headsets as part of their daily job, which requires them to be authenticated across different applications throughout the day.
- Gamers who want a seamless, hands-free and voice-free method of profile loading and authorizing in-game purchases.
- e-Commerce consumers whose check-out experience is currently interrupted by passwords.
- Surgeons, heavy machinery operators, medical professionals and others working in high-strain, sanitary environments that require especially high reliability and convenience.
- Air gapped environments where there are strict demands on performance, confidentiality and all biometric processing and decisions need to be performed on-edge devices.

To model these use-cases, we set out to perform a generalizable field test of brain-based authentication using brain signals measured noninvasively from people in their natural home environment. In other words, their real world context. This feasibility test “in the wild” advances the applied science of brain biometric analysis towards scalable implementations as all participants were completely new (naïve users) to the system and enrolled themselves from home. They used a comfortable head wearable for hours during test sessions, and this device had minimal data and battery requirements. They performed repeated authentication attempts across several different days, and individual brain ID’s were shown to be robust against changes in brain state and ambient noise inherent to brain data.

## 2. Methods

### a. Participants

Sixty-two (62) participants were recruited to complete four (4) sessions over a single (1) week at their own home. Adult participants were recruited from an opt-in screening panel and came from all five (5) major regions of the continental United States (Northeast, Southwest, West, Southeast, and Midwest). Only participants who reported normal vision, or vision that was corrected to normal with contact lenses were included. We excluded volunteers who reported using medication that might influence the experiment or other neurological or psychiatric conditions that could influence the results. Written informed consent was obtained from all participants before screening and the main experimental sessions. Thirteen (13) participants were ultimately excluded for problematic survey response patterns within the study and/or invalid brain data, leaving 49 participants (mean age= 36, SD=8.25, 16 females) enrolled and eligible to be included in the analysis.

### b. Sessions

Individuals participated in the study by recording sessions from their own homes at their own pace, over one week as detailed in Haruvi et al 2021.^27^ Each participant received an Arctop technology kit that included headphones (Sony), a brain signal measuring headband (InteraXon) and a tablet computer (Samsung) with a designated app (Arctop). Each participant recorded four sessions, one hour long each where towards the end of each session, six (6) authentication events were presented. Each authentication event started with a message declaring the upcoming event and instructing the participant to stay steady. Then, a fixation period which enabled the participant to get prepared (2 seconds) before seeing the rapid serial visual presentation (RSVP) of selected images at 10Hz for 10 seconds (Figure 2).

**Figure 2:**
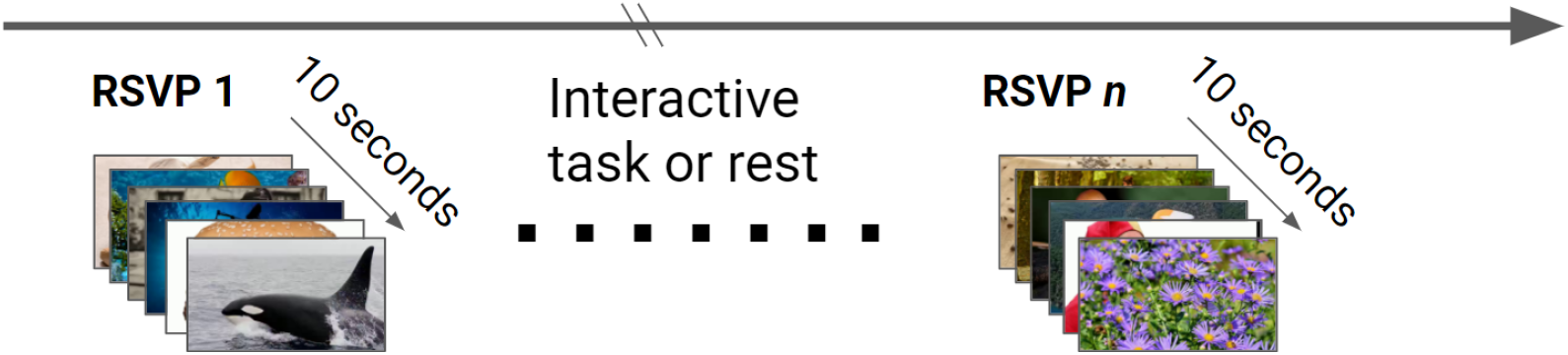
Time course of brain-based authentication using rapid serial visual presentation prompts. At each authentication event a sequence of images rapidly (10Hz) changes for 10 seconds while the brain response is recorded. In each session, six RSVP events were presented to each participant.

### c. Data Acquisition

Participants engaged in a variety of tasks during each session while their electrical brain activity was recorded using InteraXon’s Muse-S device, a portable, noninvasive electroencephalography (EEG) device weighing 41 grams (Figure 3, left panel). The device includes four dry fabric EEG sensors (sampling rate: 256 Hz), photoplethysmography (PPG) sensors (for heart rate) and motion sensors (gyroscope and accelerometer). The EEG sensors are located on the scalp, two frontal channels (AF7 and AF8) and two temporals which rest behind the ears (TP9 and TP10), with a reference channel at Fpz. The headbands were put on by the participants themselves, with the assistance of a Quality Assurance (QA) screen that started before each session. The QA showed the participants, in real-time, the channels’ quality, easily directing them to adjust the headband properly for optimal signal quality.

**Figure 3:**
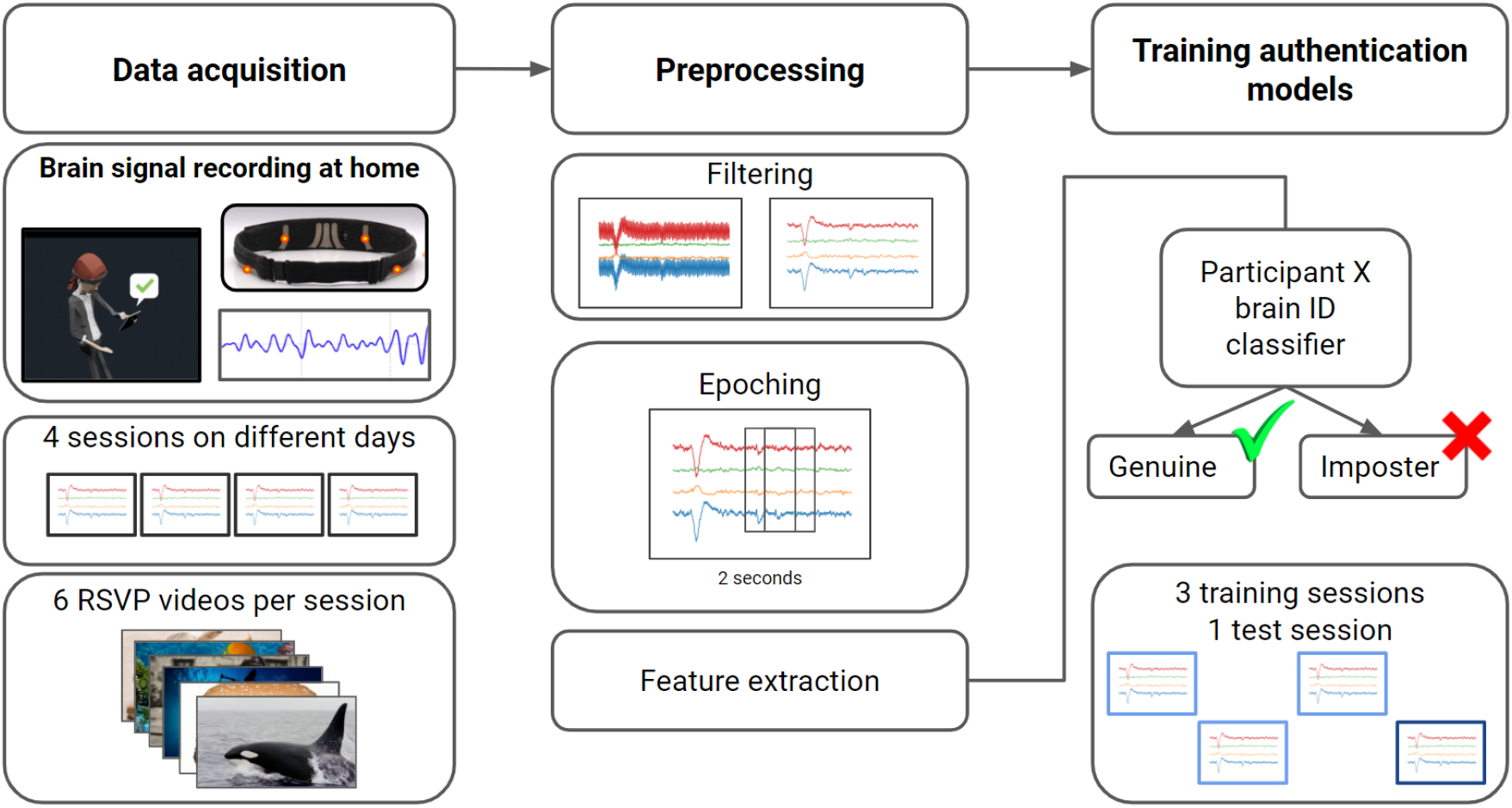
Schematic illustration of the processing pipeline. Data acquisition included at home brain data recordings of 4 sessions, each on a different day. Each session included 6 RSVP videos (Figure 2). Brain data processing included filtering the signal, feature extraction and training a machine learning authentication classifier per participant. The classifier decides if the input belongs to the participant (Genuine) or not (Imposter).

### d. Preprocessing and feature extraction

Data analysis was performed only for periods within the RSVP events. A band-pass filter (0.5-46Hz) was applied on each channel. The filtered signal of RSVP event was segmented into 31 epochs of 2 seconds in length, using a sliding window with a stride of 250ms (Figure 3, middle panels). Comprehensive feature extraction and engineering was not the goal of this current study. Here we aimed for effective information capture without deeper optimization to first test the core principles. Accordingly, for each epoch and for each EEG channel, the following features were calculated: The average power for each of the traditional frequency bands (Alpha, Beta, Gamma, Delta, Theta), power spectrum interactions (engagement index, Alpha over Delta, Beta over Theta, Theta/Alpha), time domain features such as averages, standard-deviations, kurtosis, entropy and number of zero-crossing points, and pairwise correlations between channels for the various frequency bands. All together, for each epoch, a total of 140 features were extracted.

### e. Models training and testing

For each participant we had a total of 24 RSVP events (Supp. Video 1), which we collected over the 4 sessions. For each participant, three sessions (18 events) were chosen randomly to be used for training (Figure 3, right panels). The fourth session was used for testing (6 events). Authentication prediction of an event acts in two steps, the first at the epoch level, where each epoch is determined to belong to a genuine or imposter. Second is the final decision regarding the whole event identity (genuine or imposter).

For each participant an authentication model was trained first at the epoch level. Model classification was done with XGBoost classifier (binary classification). The labeling of the data was changed in accordance to the identity under training. Epochs (feature space, 140 features per epoch) of genuine identity were labeled as one (558 epochs), while epochs from the rest of the participants were of imposter identity, and labeled as zero (26784 epochs). Thirty percent of training data (random and balance split) was dedicated for validation and to determine epoch thresholding.

Standardization procedure over the training epochs was applied. Later, standardization means and stds (standard deviation values) of the training features were used to normalize the validation and testing data. Epoch’s threshold for classification was optimized to minimize false acceptance rate (FAR), while maximizing true rejection rate (see Supp. Figure 3). Identity predictions of validation data epochs, after thresholding, exhibited high accuracy for all participants (average accuracy=0.9865, STD=0.00929).

A final decision about participant identity was given at the event level. Event threshold, as before, was determined by an optimization algorithm, but here it was done over the validation data. For the validation data, after thresholding, the average event authentication accuracy over all participants was 0.9965, with STD=0.00041. Since per each participant validation data included 264 events, it suggests that on the average, after thresholding all events were identified correctly except one.

Test data included 294 events and 9114 epochs. Training model predicted the identity probability of each epoch. Probabilities above the epoch threshold were determined to be of a genuine identity, while those below the threshold are of imposter identity (Supp. Figure 4A-C). Test event was declared to be of genuine identity only if 40% of its epochs were above the epoch threshold (Supp. Figure 4A1-C1).

## 3. Results

In our authentication system we derive from noninvasively recorded brain signals a “brain ID” abstraction that proved to be representative of each participant, and differentiating from one another. The brain response during a RSVP event is used as a brain biometric ID for identity verification. In order to demonstrate characteristics of this brain ID, we will follow the example presented in Figure 4. In our system, the authentication period depends on the event length, here it is approximately 10 seconds (one RSVP event). The event is composed of 4 channels (256Hz), segmented into 31 epochs, 2 seconds long, with a stride of 250ms (Figure 4A-B). The non-stationary nature of the brain signal, and the fact that it is a superposition of hundreds of simultaneous processes in the brain, makes the signal unique in time, unrepeatable, and unpredictable. Even when a user’s brain is stimulated by identical stimuli, no two epochs are alike (Figure 4B), nor are two events alike. In Figure 4C and in Figure 4D the epochs of two events (the brain signal space) of the same participant are presented respectively. For each event, the epochs are aligned vertically, ordered in time, creating a visualization of the brain ID. This representation enables us to see easily that none of the epochs are identical, nor are the full events.

**Figure 4:**
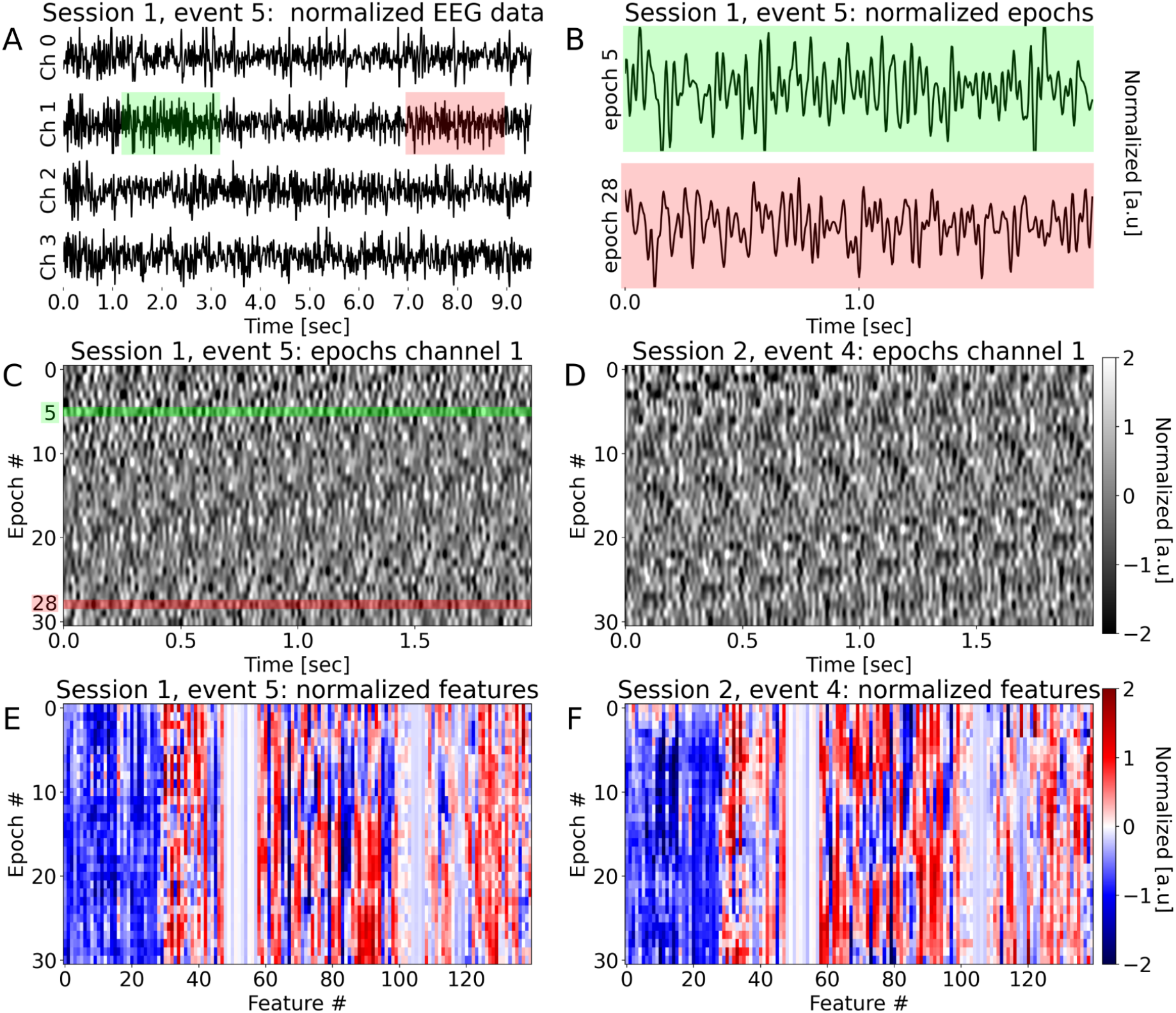
Event epoching. A. The RSVP authentication event is a normalized filtered brain signal response composed of four channels of EEG data, 10 seconds long. Each event signal is segmented into 31 epochs (where each epoch carries 4 channels), 2 seconds in length, and with a sliding window of 0.25 sec stride. B. Channel-1 of epochs #5, and #28 (top, bottom) are shown for demonstration. Note, that the shaded areas colored in green and red in panel A correspond for these epochs respectively. C. The epochs of the event signal in A, can be rearranged into an image (here again just channel-1 is shown). Where each row is an epoch, and the epochs are time ordered vertically. In C and D, events which were taken from the same participant (#39), but from different sessions are shown. E, F the corresponding features of the epochs presented in C, D are presented. Note that the calculation of epoch features involves all epoch channels. While the non-stationary nature of the EEG data dictates that the events (as shown in C, D) do not resemble each other, the features images (E, F) demonstrate high similarity.

In contrast, the brain ID data becomes highly correlated when the same events shown previously in Figure 4C-D are now presented at a higher level of analysis (the features space) (Figure 4E-F). High correlation is visible among epochs of the same event, creating a clear brain ID pattern. As one can note, a similar pattern is carried by brain data captured at different occasions, and we can conclude that usage of an event instead of a single epoch for deriving the brain ID increases the pattern robustness and increases both the sensitivity (true acceptance rate) and specificity (true rejection rate) of the system.

In Figure 5A-D, four brain IDs examples of different participants are presented. It is apparent that each brain ID carries a unique pattern that is distinguishable from the others. We would like to generalize the idea of using RSVP events brain IDs as a verification method in our authentication system over all the participants. If the event brain ID is used as an identity verification two criteria must be fulfilled:

1. The similarity between different events brain IDs of the same person is kept high: even and especially, for events which were recorded at different occasions.
2. The brain ID of each participant is unique, and distinguishable.

**Figure 5:**
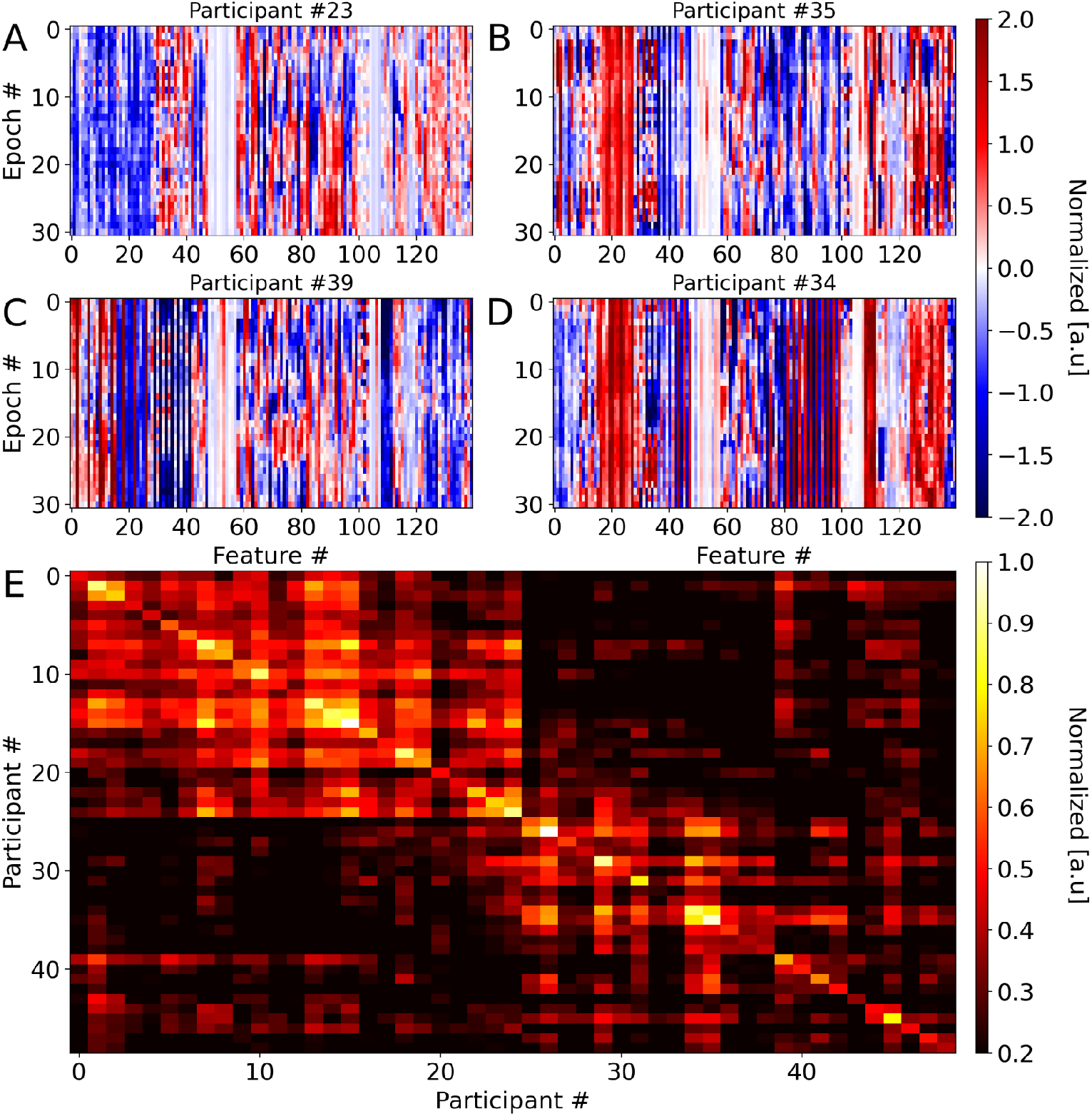
Similarity among intra and across inter participants events. Panels A, B, C, and D show the features of a single event for different participants (#23, #35, #,39, and #34 respectively). The pattern of an event appears more robust, as the features values are repeatedly conserved across many epochs. On the other hand, it looks like for each participant the pattern is specific. The similarity (or dissimilarity) between events can be measured by a correlation coefficient. In E we present the event correlation matrix, where element E*ij*, is the average pairwise correlation across all training events of participant *i* and participant *j*. Note that the intra-correlation coefficients (diagonal) are usually higher than inter-correlation (off-diagonal), suggesting that for the same participant the pattern of different events is conserved, and patterns of different participants are different. This understanding leads us to the idea of an authentication system by events. Also note that the order of the participants in E, is in accordance with the hierarchy cluster tree shown in Supp. Figure 1. The matrix here is normalized.

The similarity between two events (at the feature space) can be measured by the Pearson correlation coefficient between the means of the events. Thus the similarity between two participants is the mean of all pairwise events correlations of these participants. In Figure 5E, the normalized correlation matrix across all participants is presented. Values are represented by colors (colorbar 0.2-1), higher values suggest higher similarity. The order of participants along the axes was determined using a hierarchical clustering algorithm (see Supp. Figure 1). The averaged similarity between events belonging to the same participant (intra correlation) are along the diagonal elements of the correlation matrix, while the averaged similarity between events of two different participants (inter correlation), are the off diagonal matrix. In general, we have found that for all participants, the similarity of intra correlation is higher than the inter correlation (Figure 5E, Supp. Figure 2).

Looking more deeply, histograms in Supp. Figure 2, shows that most of the inter- and intra-participant correlation are indeed separated: for more than half of the participants the intra-correlation is higher than 0.7, where most of the inter-correlations are lower than 0.35. The inset in Supp. Figure 2 also suggests a linear relation between the mean inter-correlation of a participant and its intra correlation. Namely, participants having relatively low intra-correlation (~0.5), their inter-correlations will be low as well (~0.25). These results reflect that criteria 1 and 2 (above) are fulfilled, and the brain IDs can be used for identity classification.

As described in the Method section, for each participant an authentication model and relevant thresholds were found. These models were tested on the test data which in total included 249 genuine events, and 14112 imposter events. The general performance of our authentication system is summarized in Figure 6. The averaged false acceptance rate (FAR) is 9% and the false rejection rate (FRR) is 13%, making the solution sufficient for certain commercial authentication use-cases, but not all. The averages shown here are the means over the individuals’ FAR, FRR.

**Figure 6:**
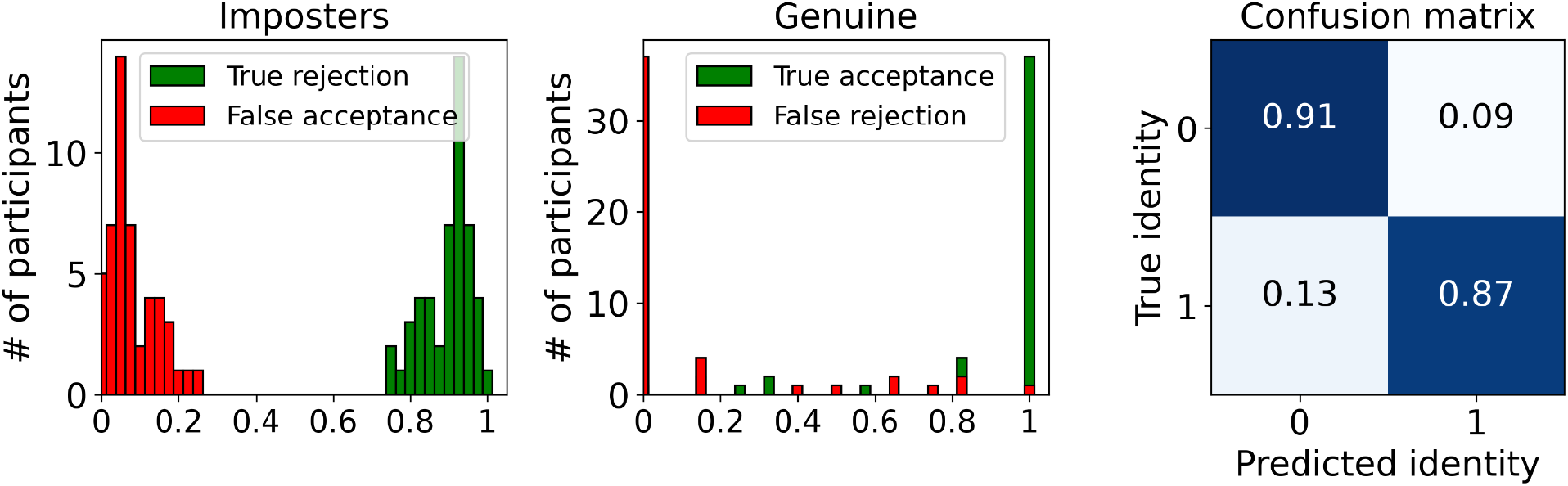
Summary of authentication performance in the field. Forty nine participants were included in the test. Each participant had six genuine events, and 288 imposter events. All together in this test we used 294 genuine events, and 14112 imposter events. In A, and B histograms of authentication system performance at the participant level is present. In A, the performance regarding imposters (true rejection rate, and false acceptance rate). In B, the performance regarding genuines identities (true acceptance rate, and false rejection rate). These values were first calculated per each participant, and then distribution was calculated. C. A confusion matrix summary, showing the averaged performance over all participants. A detailed performance summary per each participant can be found in supp. Table 1.

Out of the total number of participants in this experiment (49), 37 participants have FRR=0, where 24 participants have FRR=0 and FAR<=9% (Supp. Table 1).

Next we asked what will be the performance of the authentication system when only certain brain signal information is considered. Explicitly, we repeated the training procedure (Methods) but this time with only the power spectrum features of the following brainwave modes: Delta(0.5-4Hz), Theta(4-8Hz), Alpha(8-12.5), Beta(12.5-30Hz) and Gamma(30-48), and with some combinations (Alpha-Beta, and Theta-Alpha-Beta). We found that usually for these features, FRR can reach low values while the FAR is always kept high (Table.1). As the number of features is increasing, the better the performances of the authentication system. This implies that more sophisticated models such as deep neural networks will greatly improve the performance of the authentication system. We will report on the results of different systems such as these in future field test reports.

**Table 1:** Model performance as function of feature types. The same training and testing datasets were used for all models. The power spectrum density (PSD) of the following frequency bands were used as features. Delta(0.5-4Hz), Theta(4-8Hz), Alpha(8-12.5), Beta(12.5-30Hz) and Gamma(30-48). Each bandpass has four features, corresponding to the number of brain data channels. When using only one type of powerband feature, the averaged FRR may reach low levels, but the FAR always remains high.

| Features type | False acceptance rate | False rejection rate |
| --- | --- | --- |
| Delta | 0.83 | 0.02 |
| Theta | 0.82 | 0.02 |
| Alpha | 0.75 | 0.03 |
| Beta | 0.36 | 0.17 |
| Gamma | 0.28 | 0.22 |
| Alpha Beta | 0.30 | 0.14 |
| Theta Alpha Beta | 0.27 | 0.13 |
| All PSD brain waves | 0.17 | 0.16 |
| All features | 0.09 | 0.13 |

## 4. Discussion

We performed a generalizable field test of a brain-based authentication system that uses noninvasively measured brain signals to verify user identity. All participants were completely new (naïve users) to the system, enrolled themselves from home in a self-guided tutorial, used a comfortable head wearable for hours at a time without issue and performed repeated authentication measures across multiple days. On the whole, this amounts to a reasonable simulation of real contexts that enterprise and consumer authentication methods need to operate in to be commercially viable. Specifically, these methods must work regardless of time of day and be robust to changes in brain state (pre/post-coffee, hunger, wakefulness, awareness, etc.) and the ambient noise inherent to measurements made outside of controlled laboratory conditions.

The main goal of this field test was to evaluate the base feasibility of a scalable, commercial-grade brain ID authentication system; advanced data engineering methods were not applied to boost performance further. A simplified feature set and simple machine learning methods were applied over a minimal training period of less than three minutes enrollment data per participant. Amidst these severe constraints on performance, brain-based authentication proved to be approaching commercial-grade levels. In future tests the parameters used will be optimized, here our authentication system ran on suboptimal parameters that were fast calculated to serve as more heuristics than anything. For example the epoch length, the authentication event duration, selection of features by their importance, or the amount of training data we know have an impact on performance from previous and ongoing work. These are all tunable parameters depending on the demands of the authentication task: future research will clarify the timescales at which the optimal information for identification verification occurs for each tier of authentication system. We are confident that more sophisticated machine-learning architectures together with other parameters optimization will deliver superior, product level authentication performance that will match or exceed the performance of top non-brain authentication biometrics available today.

Wearables that touch the head, such as headphones or AR/VR, are a natural form factor for brain-based authentication and we anticipate that demand from enterprises and consumers will necessitate that these devices evolve beyond passwords and fingerprints to iris ID based on eye scanning and eventually brain ID, based on brain scanning. The demand for both strong and convenient authentication solutions for future headworn devices drove our design of the paradigm for prompt-response analysis here, and it is notable that the rapid image prompt-response paradigm evaluated (with users watching images on a tablet while wearing a headband) has been validated by us elsewhere in AR (Supp. Video 2) using Microsoft Hololens.

Given the performance obtained in this field test and the ease-of-use of this method for head wearables, brain ID seems to be one of the most intuitive and powerful authentication solutions for next generation headworn computers. Brain identities, like any other biometric identity, will need to conform to privacy standards and be offered within protected software and chip architectures such as those pioneered for fingerprint scanners and face recognition, but this is no limitation on the adoption of such a beneficial method.

Biometrics as a class are uniquely comfortable and convenient to use because they do not require the user to remember anything (like a password), or carry anything (like a physical key). Biometrics offset the cognitive load of password management plus the risks associated with alphanumeric passwords, and even offer the promise of obviating passwords altogether in future computing ecosystems. For now, brain ID is at a nascent stage of industry adoption and the solution presented here represents one of the more scalable designs, since we can easily increase the dataset to more participants and more events within the principled framework of forcing divergences in inter-participant data and convergences in intra-participant data.

Furthermore, the head wearable that people put on themselves in this test to measure their brain signal is a consumer device that is currently available worldwide, highlighting the lack of need for exotic or rare materials to acquire sufficient brain signal to measure brain IDs, nor the need for specialized laboratories or facilities. More information in the brain signal remains unexplored here, being outside the scope of the current field test and report. Future research will develop concepts related to the theoretical and practical information boundaries in brain signal, since for head wearables in particular, brain biometric identity warrants continued testing across expanded participant populations and implementation in commercial devices that are optimized for given use-cases and environments.

## 5. Conclusion

We showed that a relatively simple brain-based authentication system can use noninvasively measured brain signals from consumer quality head wearable devices to differentiate between users with a high degree of certainty. Authentication using noninvasively-measured brain signals in this way was found to not only be feasible, but robust: the correlation matrices derived from the current test find our computed brain identities to be readily distinguishable between different participants and consistently similar among participants, satisfying the core requirements of a commercial-grade biometric authentication system. The complexity inherent to human brain signals was, therefore, found to not be too volatile to be leveraged for steady, reliable use as a passwordless authentication method.

We built and validated the method through a scalable software infrastructure that was able to deliver brain-based authentication at a commercial-grade, within a generalized framework that provides for continual performance improvement with additions of new participants. As the methods were designed around characteristic patterns observable during limited windows of time, at any time, it is clear that there is value to continued data collection at larger scales and across additional contexts. For both inter-subject variability and to further clarify the invariant patterns underlying intra-participant variability, expanded data collection can be beneficial. The present sample is sufficient however to conclude that brain-based authentication is already a viable method for certain commercial uses, and has the potential to serve many more in the future.

## Author information

Affiliations

Arctop Inc., R&D, Kaufmann St. 4, Tel Aviv-Yafo, 6801296, Israel.

## Ethics declarations

All authors are employees of Arctop Inc.

## SUPPLEMENTARY MATERIALS

**Supp. Figure 1:**
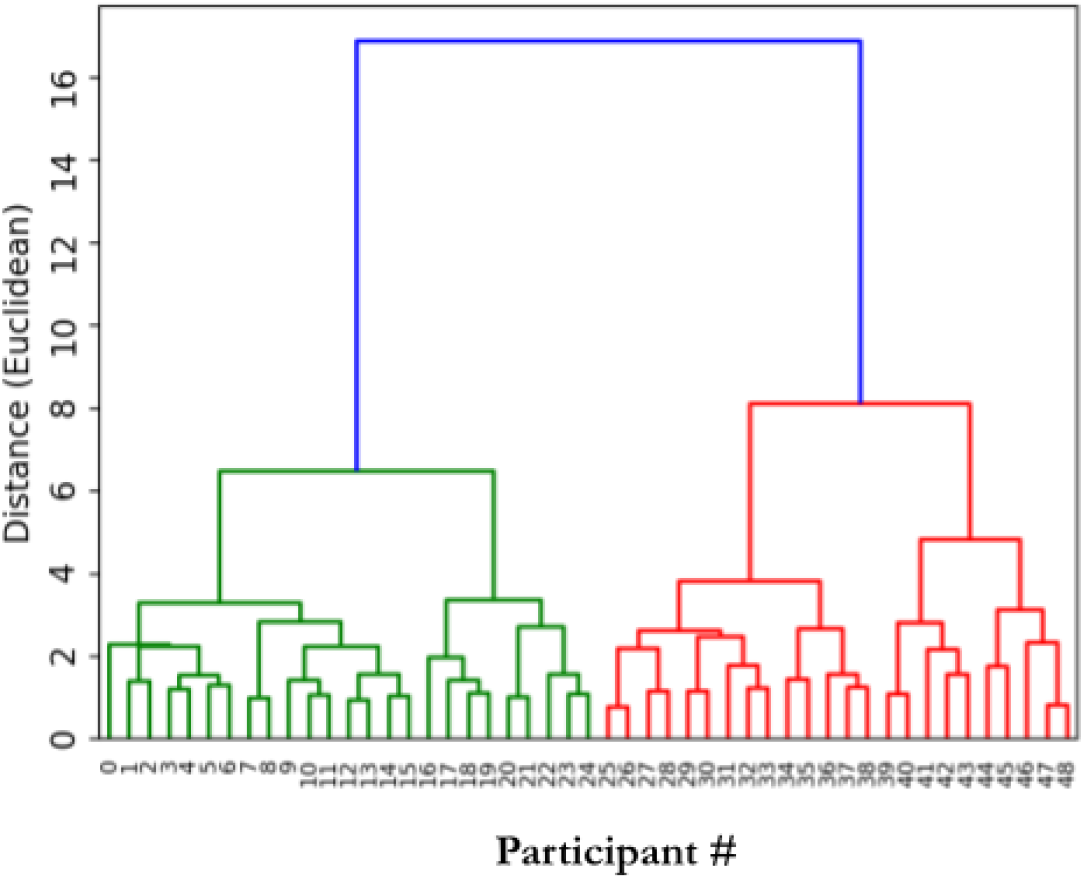
Brain biometric ID cluster tree. The mean overall training events were calculated for each participant. Mean event correlation matrix between participants was then calculated by pairwise correlation. Using this matrix, the hierarchical cluster tree (dendrogram) algorithm creates the linkage distance between participants (y-axis).

**Supp. Figure 2:**
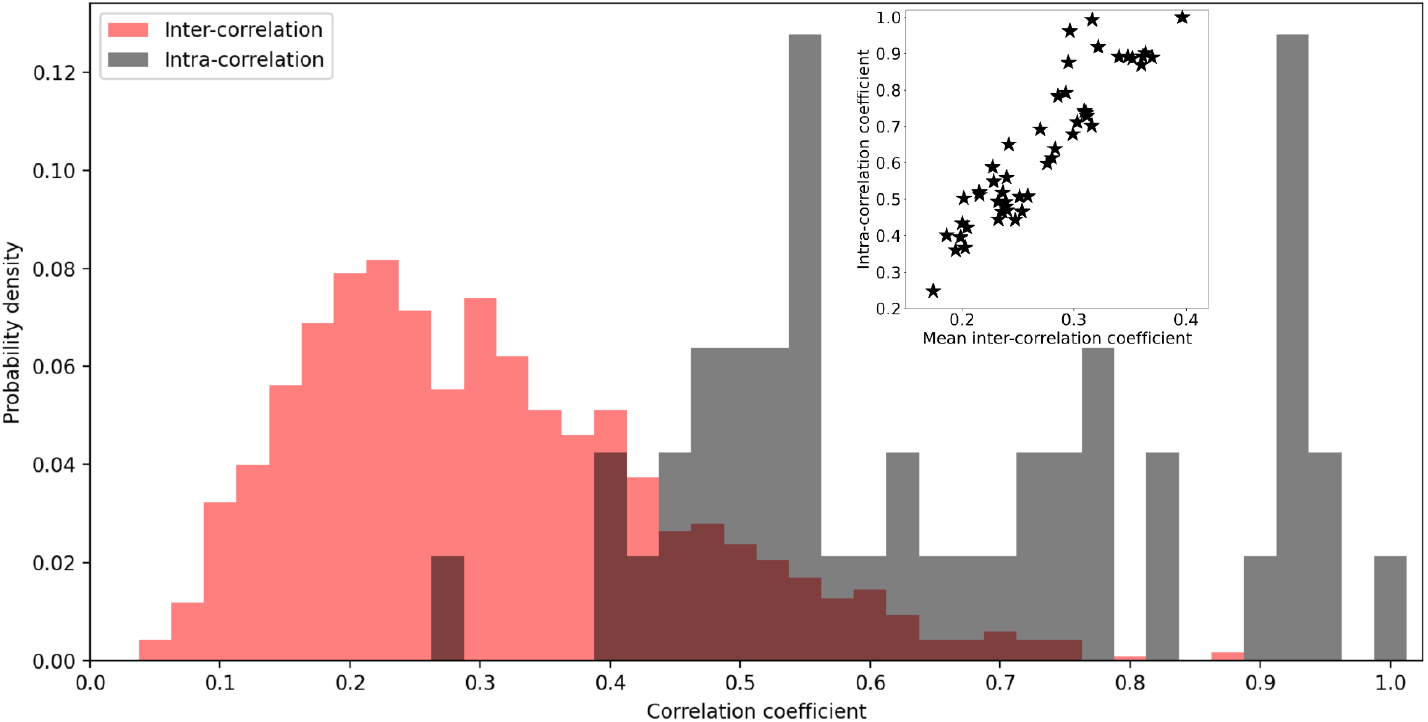
Histograms of intra-participant and inter-participants events correlations. Intra-participant events correlation is the mean of pairwise correlation between all training events of a participant with themselves. Inter-participant correlations are the mean of pairwise correlation of all training events of a participant with the events of another participant. The diagonal elements in the event correlation matrix (Figure 6E), represents the intra-participant correlations while the inter-participants correlations are the off-diagonal elements of the matrix. It is clearly seen that intra-participant correlations are generally higher than the inter-participants correlation. Meaning, a higher similarity within intra events patterns compared with inter-participants events. Although there is an overlap between the two histograms, it does not necessarily mean that perfect separability at the authentication event level is not feasible, as suggested by the inset.

**Supp. Figure 3:**
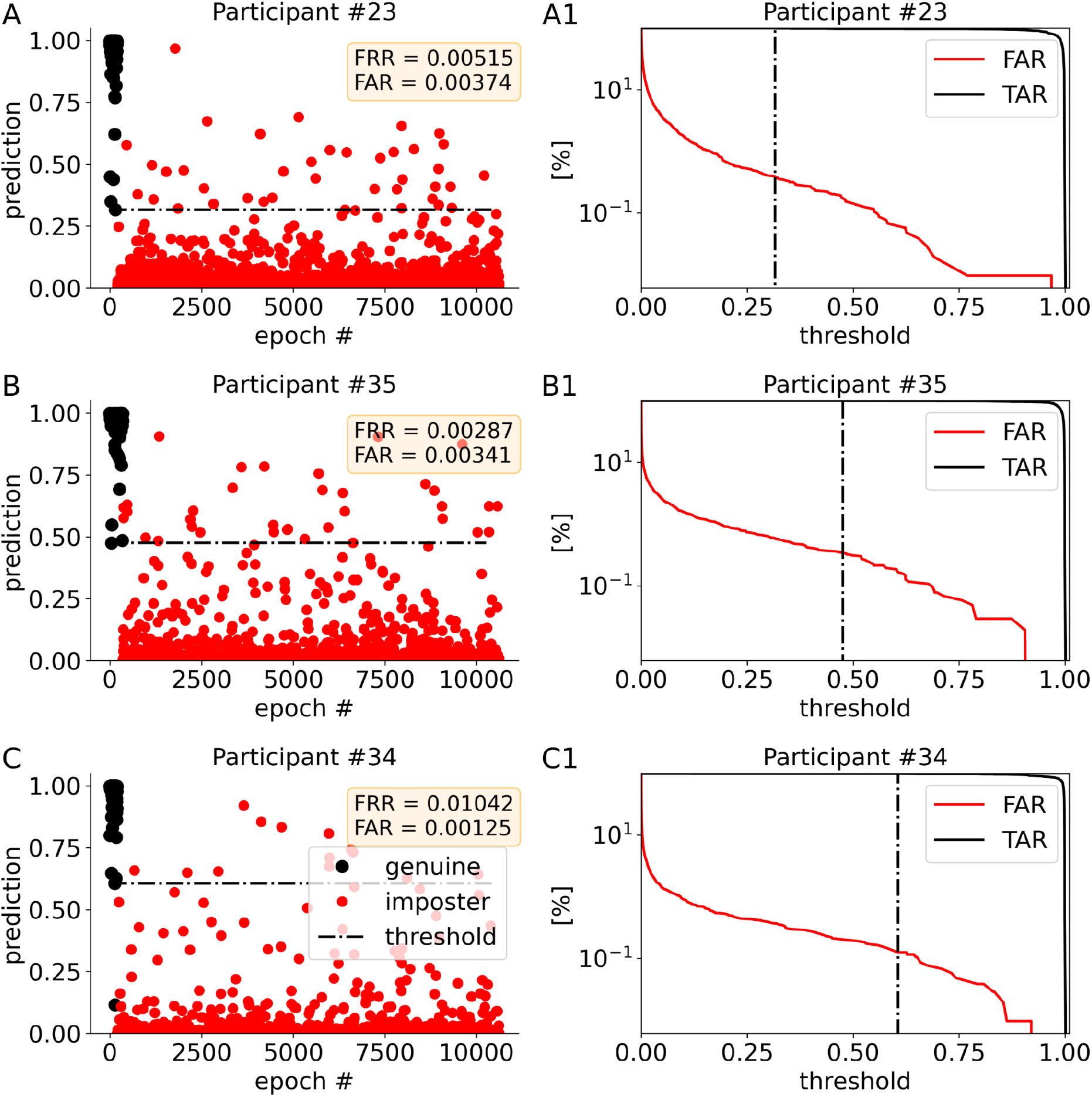
Epoch predictions and threshold determination. Three sessions per participant are contributing to the training data. Out of it, 30% are devoted for model validation, and to determine the model threshold. In panels A, B, C the prediction of three models trained for three participants (sub #23, # 35, #34 respectively) are presented for the validation data. Here, epochs predictions of genuine identity are marked in black dots, and epochs predictions of imposters are marked in red. The threshold (black dashed line), discernmenting between genuine and imposter epochs is determined by an optimization algorithm. The algorithm finds a threshold probability in which the false acceptance rate (FAR) is minimal while the true acceptance rate (1-FRR) is maximal. This is under the condition for TAR>90%, and FAR<3%. This is demonstrated in panels A1, B1, C1. FAR, TAR functions are plotted in red and black respectively, the threshold which was found is marked in black dashed-dot line, y-axis is in logarithmic scale.

**Supp. Figure 4:**
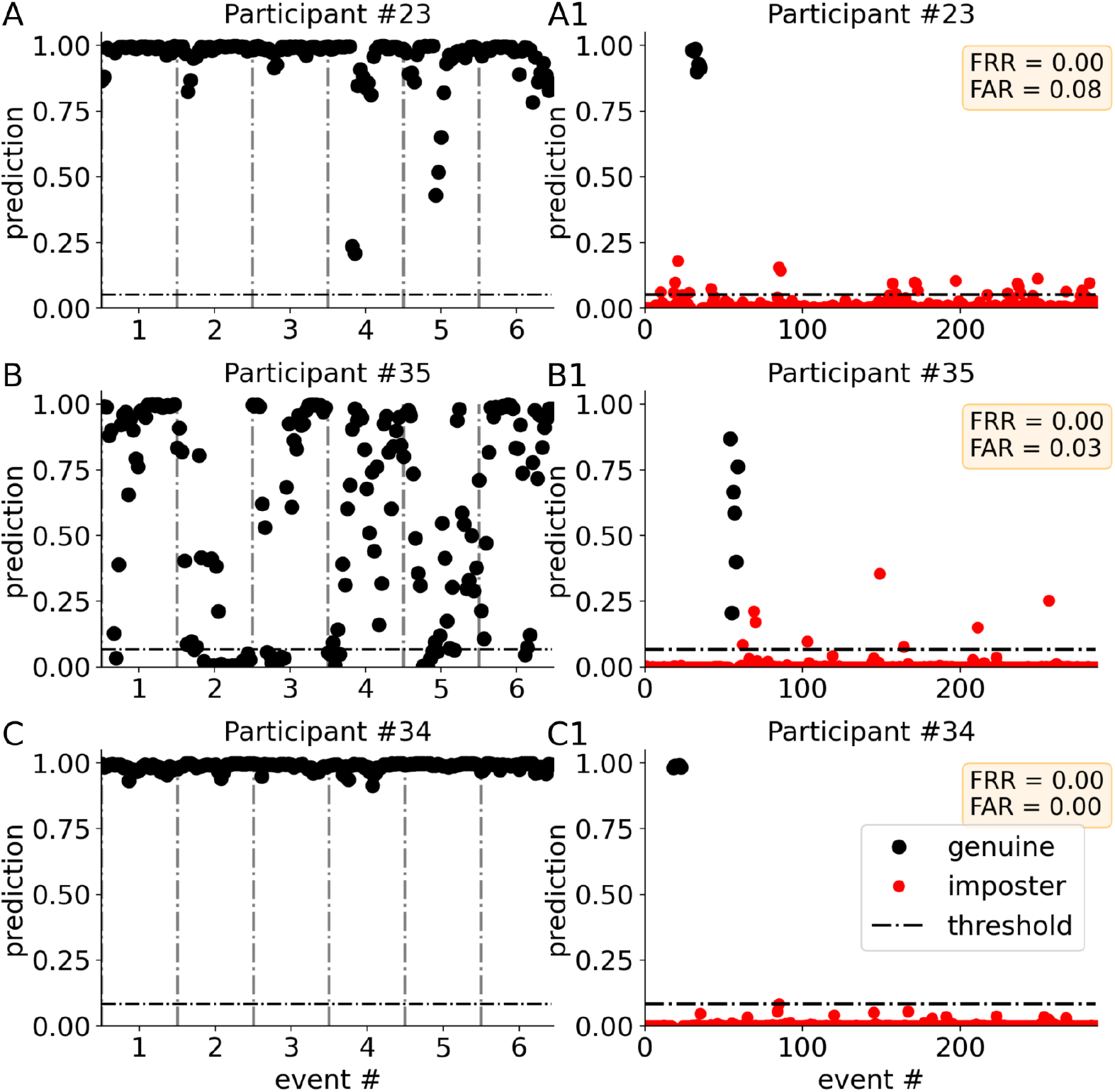
Event prediction. The events are segmented into 31 epochs. The probability of each epoch to be of a genuine identity or of an imposter one is determined by a model. Probabilities above the threshold (black dash line) belong to genuine identity, and if below the threshold, to an imposter. Threshold was determined previously in the training process (see Supp. Figure 3). A, B, C the predictions of three models trained for three participants (sub #23, # 35, #34 respectively) are presented. Here the epochs under test are only of genuine identity (black dots). While in A, and C all predictions are above threshold, in B some of the predictions are below the threshold. The final decision whether the event is of genuine identity is determined only if more than 40% of epochs are above the threshold. In A1, B1, C1 full test prediction is shown for the same three participants. The test included 294 events, where each participant has 6 genuine events. Events of genuine identity are marked in black dots and imposter events are in red dots. In all three cases all genuine events were identified correctly, having zero false rejection rate (FRR=0). As for the imposters, only in C1, all imposter events are below the threshold, with zero false acceptance rate (FAR=0). The final FRR and FAR of each participant is shown in the yellow windows.

**Supp. Table 1:** A detailed performance of the authentication system for each participant. The coefficients of the confusion matrix per each participant is presented.

| Subject # | True rejection rate (TRR) | True acceptance rate (TAR) | False acceptance rate (FAR) | False rejection rate (FRR) |
| --- | --- | --- | --- | --- |
| 0 | 0.81 | 0.83 | 0.19 | 0.17 |
| 1 | 0.95 | 1.00 | 0.05 | 0.00 |
| 2 | 0.82 | 1.00 | 0.18 | 0.00 |
| 3 | 0.93 | 1.00 | 0.07 | 0.00 |
| 4 | 0.86 | 1.00 | 0.14 | 0.00 |
| 5 | 0.86 | 1.00 | 0.14 | 0.00 |
| 6 | 0.87 | 0.60 | 0.13 | 0.40 |
| 7 | 0.95 | 1.00 | 0.05 | 0.00 |
| 8 | 0.85 | 1.00 | 0.15 | 0.00 |
| 9 | 0.78 | 1.00 | 0.22 | 0.00 |
| 10 | 0.92 | 1.00 | 0.08 | 0.00 |
| 11 | 0.92 | 1.00 | 0.08 | 0.00 |
| 12 | 0.76 | 1.00 | 0.24 | 0.00 |
| 13 | 0.84 | 0.83 | 0.16 | 0.17 |
| 14 | 0.97 | 1.00 | 0.03 | 0.00 |
| 15 | 0.95 | 0.83 | 0.05 | 0.17 |
| 16 | 0.92 | 1.00 | 0.08 | 0.00 |
| 17 | 0.94 | 0.17 | 0.06 | 0.83 |
| 18 | 0.96 | 1.00 | 0.04 | 0.00 |
| 19 | 0.91 | 1.00 | 0.09 | 0.00 |
| 20 | 0.90 | 1.00 | 0.10 | 0.00 |
| 21 | 0.89 | 1.00 | 0.11 | 0.00 |
| 22 | 0.84 | 0.33 | 0.16 | 0.67 |
| 23 | 0.99 | 1.00 | 0.01 | 0.00 |
| 24 | 0.96 | 1.00 | 0.04 | 0.00 |
| 25 | 0.91 | 1.00 | 0.09 | 0.00 |
| 26 | 0.99 | 0.50 | 0.01 | 0.50 |
| 27 | 0.93 | 1.00 | 0.07 | 0.00 |
| 28 | 0.95 | 1.00 | 0.05 | 0.00 |
| 29 | 0.97 | 1.00 | 0.03 | 0.00 |
| 30 | 0.81 | 1.00 | 0.19 | 0.00 |
| 31 | 0.94 | 1.00 | 0.06 | 0.00 |
| 32 | 0.75 | 1.00 | 0.25 | 0.00 |
| 33 | 0.93 | 0.33 | 0.07 | 0.67 |
| 34 | 1.00 | 1.00 | 0.00 | 0.00 |
| 35 | 0.98 | 1.00 | 0.02 | 0.00 |
| 36 | 0.94 | 1.00 | 0.06 | 0.00 |
| 37 | 0.94 | 0.83 | 0.06 | 0.17 |
| 38 | 0.96 | 1.00 | 0.04 | 0.00 |
| 39 | 0.95 | 1.00 | 0.05 | 0.00 |
| 40 | 0.85 | 1.00 | 0.15 | 0.00 |
| 41 | 0.94 | 0.00 | 0.06 | 1.00 |
| 42 | 0.96 | 1.00 | 0.04 | 0.00 |
| 43 | 0.92 | 1.00 | 0.08 | 0.00 |
| 44 | 0.86 | 1.00 | 0.14 | 0.00 |
| 45 | 0.98 | 0.17 | 0.02 | 0.83 |
| 46 | 0.95 | 0.25 | 0.05 | 0.75 |
| 47 | 0.89 | 1.00 | 0.11 | 0.00 |
| 48 | 0.96 | 1.00 | 0.04 | 0.00 |
| Average | 0.91 | 0.87 | 0.09 | 0.13 |

**Supp. Video 1: Rapid Serial Visual Presentation (RSVP) Stimuli.** Example of a stream of images watched by participants while brain signals were recorded by their headband. https://youtu.be/TWUzbX3Q8sk

**Supp. Video 2: Brain-based Authentication: Living Room Demo.** Microsoft HoloLens 1, retrofitted with BCI sensors, delivers passwordless authentication. https://youtu.be/n6v9z3lNs2M

